## Supplemental Figure legends for "Mitochondrial translocation of TFEB regulates complex I and inflammation"

**Supplementary Figure Legends**

**S1. (A)** Volcano plot of FLAG-immunoprecipitates from cells transfected with WT-TFEB FLAG and an unrelated protein as control. The blue dots, green dots and red dots represent positive interactors, established TFEB-interactors and high enrichment of TFEB respectively, indicating the reliability of the proteomics data. **(B)** Boxplot of 14-3-3 proteins in the core TFEB interactome. Data represent the means ± S.E.M. **(C)** TFEB and GAPDH protein and **(D)** *TFEB* mRNA expression (normalized to HPRT) in control (shCTRL) and TFEB-depleted (shTFEB) HeLa cells. Western blot and qPCR are representative of three independent experiments showing similar results. Shown are mean ± SEM, n=3 biological replicates. **(E)** LC3B and endogenous TFEB levels in shCTRL and shTFEB cells in Untreated conditions (UT) and upon torin-1 treatment for 1 hour (1 h). The experiment was repeated independently three times showing similar results. **(F)** TFEB localization in subcellular fractions isolated from human monocyte derived macrophages. LAMINB, TOMM20, and GAPDH served as controls for nucleus, mitochondria, and cytoplasm, respectively. Western blot is representative of two independent experiments showing similar results.

**S2. (A)** mtDNA and TOMM20 staining in HeLa cells to confirm mitochondrial localization of mtDNA. Scale bar = 10 $\mu$m. **(B)** Protease protection assay on mitochondria isolated from HEK293T cells after treatment with increasing concentrations of trypsin (0 – 75 $\mu$g) in the presence or absence of swelling buffer. Samples were analyzed for the outer mitochondrial membrane (OMM) protein TOMM20, the inner mitochondrial membrane (IMM), protein TIM23, and the mitochondrial matrix (MM) protein MRPL12. MRPL12 and endogenous TFEB were exposed to trypsin after mitochondria were lysed with Triton X-100. The experiment was repeated twice with similar results. **(D)** Densitometric analysis of protein levels in Supplementary Fig. 2B. Band intensities were normalized to the first lane (untreated mitochondria) of the respective protein.

**S3.** (**A)**, Analysis of subcellular fractions isolated from cells expressing FLAG alone, FLAG-tagged WT-TFEB or FLAG-tagged ΔMTS-TFEB. LAMINB, TOMM20 and GAPDH served as controls for nucleus, mitochondria and cytosol, respectively. Relative abundance of TFEB in the various fractions. The experiment was repeated three times with similar results. **(B)** Relative abundance of TFEB in the various fractions of the immunoblot shown in Figure S3A. **(C)** Relative abundance of TFEB in the various fractions of the immunoblot shown in Figure 3B and another two biological replicates. **(D)** HeLa cells stained for endogenous TOMM20, TFEB, and nucleus. Scale bar = 10 μM. 25 cells per group were analyzed. **(E)** Colocalization was evaluated in ROIs (25x25 pixels) as described in Methods section. Mander's colocalization coefficients using the calculated thresholds (tM) were determined for the green (tM1) (TFEB) and the red (tM2) (TOMM20) channels. **(F)** TOMM20 staining in HeLa cells transfected with FLAG-tagged WT-TFEB or FLAG-tagged MLS-TFEB. Scale bar= 10 μm. Ten cells per group were analyzed. Colocalization was evaluated in ROIs (25x25 pixels) as described in Methods section. Red bars show the Mander's colocalization coefficients using the calculated thresholds (tM) for the green (FLAG) and the red (TOMM20) channels. Blue bars show the Mander's colocalization coefficients using the calculated thresholds (tM) for the green (FLAG) and the blue (DAPI) channels. **(G)** HLH-30::mNeonGreen expressed in muscle and hypodermal cells of *C. elegans* (wild type, day 1 adult) and mitochondria stained with MitoTracker Deep Red. Scale bar = 10 μm.

**S4. (A)** Percentage of perinuclear and cytosolic mitochondria clustering. n=50 cells each. **(B)** Basal respiration rate, ATP-linked respiration and Maximal respiration in HEK293T transiently transfected with control siRNA (siCTRL), siRNA against TFEB (siTFEB). Shown are mean ± SEM, n=3 biological replicates. **(C)** Schematic representation of the ΔNLS-TFEB FLAG plasmid in which the basic residues (R245–R248) were mutated to alanine, to prevent TFEB nuclear translocation; and the MLS motif was mutated to prevent TFEB mitochondrial translocation; S142A/S211A-TFEB FLAG plasmid (in which the Ser142 and Ser211 residues were mutated to alanine, to prevent TFEB cytosolic retention; and the MLS motif was mutated to prevent TFEB mitochondrial translocation). **(D)** Western Blot of subcellular fractions isolated from cells expressing FLAG-tagged WT-TFEB or FLAG-tagged ΔNLS-TFEB or FLAG-tagged S142A/S211A-TFEB. LAMINB, TOMM20 and GAPDH served as controls for nucleus, mitochondria and cytoplasm, respectively. The experiment was repeated two times with similar results. **(E)** HeLa cells transfected with FLAG-tagged ΔNLS-TFEB and FLAG-tagged S142A/S211A-TFEB were stained using anti-FLAG. Scale bar= 10 μm. **(F)** OXPHOS profile normalized to Basal respiration of shCTRL cells and shTFEB cells transfected with FLAG, WT-TFEB FLAG, MLS-TFEB FLAG, ΔNLS-TFEB FLAG and S142A/S211A-TFEB FLAG. **(G)** Basal respiration rate, ATP-linked respiration and Maximal respiration normalized to Basal respiration, in shCTRL and shTFEB cells transfected with FLAG, WT-TFEB FLAG, MLS-TFEB FLAG, ΔNLS-TFEB FLAG and S142A/S211A-TFEB FLAG. Shown are mean ± SEM, n=3 biological replicates.

**S5. (A)** 1D enrichment of the whole proteome analysis in shCTRL versus shTFEB cells. Proteins were annotated by the Gene Ontology term. **(B)** $log_2$ fold change of all detected proteins and proteins associated with Complex I in shCTRL and shTFEB-depleted cells. **(C)** Boxplot and individual points represent proteins of specific complex I modules**. (D)** ATP5a, ATP5b, COX2, UQCRC2, SDHA, SDHB, TFEB, and SLP2 expression in shCTRL and shTFEB cells. (**F)** Mean densitometric analysis of Western blot shown in Figure S5D. Data shown is representative of two independent experiments showing similar results.

**S6. (A)** Immunoblot of TFEB and GAPDH upon treatment with the LONP1 inhibitor CDDO. **(B)** Mean densitometric analysis of the immunoblot from S6A. Shown are mean ± SEM. The experiment was repeated three times with similar results. **(C)** Mean densitometric analysis of the immunoblot from Figure 6C. Shown are mean ± SEM. The experiment was repeated two times with similar results. **(D)** mRNA expression of LONP1 (relative to HPRT) in shCTRL and shTFEB cells. Shown are mean ± SEM, n=3 biological replicates. **(E)** mRNA expression of TFEB, LONP1, NDUFA8, NDUFB10, mtND6, NDUFS1 and NDUFS5 (relative to RPL13A) in shCTRL and shTFEB cells transfected with siCTRL and siLONP1. Shown are mean ± SEM, n=3 biological replicates with 6 technical replicates. **(F)** SyproRuby staining of SDS-PAGE depicting the purity of LONP1 and TFEB protein isolation in Figure 6I.

**S7. (A)** Endogenous TFEB localization in uninfected cells (UI) *S.* Typhimurium-infected (*S*.T.) and torin-1 (30 minutes) treated HeLa cells. Scale Bar = 4 $\mu$m. **(B)** HeLa cells from Figure S1A were analyzed to calculate the percentage of cells showing nuclear TFEB localization. Shown are mean ± SEM, n=15 images. The experiment was repeated independently three times with similar results. **(C)** Relative mRNA expression of IL-6, IL-8 and IL-1α (relative to RPL13A) in shCTRL and shTFEB cells. Shown are mean ± SEM, n=4 biological replicates. **(D)** WT-TFEB-FLAG subcellular localization upon 24 h of infection with *S.* Typhimurium infection (MOI 100) in HeLa cells. LAMINB, TOMM20, and GAPDH served as controls for nucleus, mitochondria, and cytosol, respectively. Western blot is representative of two independent experiments showing similar results. **(E)** Relative mRNA levels of IL-6 and IL-8 (relative to RLP13A) in shCTRL and shTFEB transfected with FLAG, FLAG-tagged WT-TFEB, ΔNLS-TFEB or S142A/S211A-TFEB plasmids and infected for 6 h with *S.* Typhimurium (MOI 100). Shown are mean ± SEM, n=3 biological replicates. **(F)** IL-6 expression in supernatants of shCTRL and shTFEB cells untreated (UT) and treated with 2.5 μM of CDDO infected for 24 h with *S.* Typhimurium (MOI 100). Shown are mean ± SEM, n=3 biological replicates. **(G)** IL-6 in supernatants of shCTRL and shTFEB cells untreated (UT) or treated with 500 nM of Rotenone (Rot) infected for 24 h with *S.* Typhimurium. Shown are mean ± SEM, n=3 biological replicates. **(H)** MitoSOX-based flow cytometric detection of mitochondrial ROS production in shCTRL and shTFEB cells untreated and treated with CDDO. Shown is the Mean Fluorescence Intensity (MFI). Shown are mean ± SEM, n=3 biological replicates.
